# Characterization of pathology-inducing α-synuclein species from human diseased brain tissue

**DOI:** 10.1101/588335

**Authors:** John D. Graef, Nina Hoque, Craig Polson, Ling Yang, Lawrence Iben, Yang Cao, Nino Devidze, Michael K. Ahlijanian, Jere E. Meredith

**Author notes:** **Corresponding Author** Michael K Ahlijanian, Ph.D.

## Abstract

Synucleinopathies are a group of neurodegenerative diseases characterized by the presence of pathological accumulations of misfolded, phosphorylated α-synuclein (αSyn) protein. Multiple lines of evidence indicate that synucleinopathy disease progression is driven by a prion-like process of transmission of a pathologic form of αSyn. One potential therapeutic approach to prevent cell-to-cell propagation is to target this transmissible species with selective antibodies. In this study, a rodent primary neuronal culture reporter system was developed to monitor induction of detergent-insoluble, phosphorylated (pS129) aggregates of αSyn. Induction of pS129 αSyn pathology was observed with both synthetic αSyn fibrils (PFFs) and brain lysates from multiple system atrophy (MSA) patients but not αSyn monomers or human brain lysate controls. The induction-competent species in MSA lysates could be enriched by high-speed centrifugation suggesting that it is present as a high molecular weight aggregate. Furthermore, samples derived from brain lysates from Parkinson’s disease (PD) and Dementia with Lewy Bodies (DLB) patients also induced pS129 αSyn pathology, but required longer incubation times. Lastly, the potential of αSyn selective antibodies to immunodeplete induction-competent forms of αSyn from both PFF and synucleinopathy brain samples is described. The results demonstrate that antibodies targeting the C-terminal of αSyn are most effective for immunodepletion of pathology-inducing forms of αSyn from samples derived from human synucleinopathy brains. Furthermore, the data support the hypothesis that antibodies that recognize a C-terminal epitope and exhibit selectivity for oligomeric forms over monomeric forms of αSyn represent a desirable target for immunotherapy for synucleinopathy patients.

## Introduction

α-synuclein (αSyn) is a 140 amino acid protein preferentially expressed in neurons at pre-synaptic terminals where it is thought to play a role in regulating synaptic transmission (1). It has been proposed to exist natively as both an unfolded monomer (2) and a stable tetramer of α-helices (3, 4) and has been shown to undergo several posttranslational modifications (5). One modification that has been extensively studied is phosphorylation of αSyn at amino acid serine 129 (pS129). Normally, only a small percentage of αSyn is constitutively phosphorylated at S129, whereas the amount of pS129-positive αSyn present in pathologic intracellular inclusions is vastly increased (6). These pathological inclusions consist of aggregated, insoluble accumulations of misfolded αSyn proteins and are a characteristic feature of a group of neurodegenerative diseases collectively known as synucleinopathies (7).

The pathological aggregates of αSyn found in neurons are called Lewy bodies and are the characteristic hallmarks of both Parkinson’s Disease (PD) and dementia with Lewy bodies (DLB). Additionally, abnormal αSyn-rich lesions called glial cytoplasmic inclusions (GCIs) are found in oligodendrocytes, and represent the hallmark of a rapidly progressing, fatal synucleinopathy known as multiple systems atrophy (MSA). The idea that the progression of synucleinopathies is in part due to cell to cell transmission of pathologic forms of αSyn species has gained support in recent years (8–10). The initial evidence for this hypothesis derives from the observed stereotypical progression of brain pathology described in PD (11) and from evidence of host-to-graft spreading of αSyn aggregates in PD patients (12). In addition, reports of either undetectable (13–15) or low levels (16) of αSyn mRNA expression in oligodendrocytes suggests that a pathological form of αSyn is propagated from neurons to oligodendrocytes in MSA. Recent work supports this idea of αSyn propagation, demonstrating that αSyn is taken up by oligodendrocytes both *in vitro* and *in vivo* (17). Moreover, inoculation of human brain homogenates from MSA patients into αSyn over-expressing transgenic mice results in neurological dysfunction and extensive pS129-positive neuronal deposits, supporting the hypothesis that MSA may be transmitted by the prion-like propagation of pathologic forms of αSyn (10, 18).

In this study, an *in vitro* reporter assay was developed using rat hippocampal neuronal cultures overexpressing the human A53T αSyn mutant protein. The A53T mutation is associated with familial forms of PD (19) and has been shown to accelerate αSyn fibrillization *in vitro* (20) leading to greater neurotoxicity in transgenic mice (21). We used both synthetic, pre-formed αSyn fibrils (PFFs) and brain lysates from MSA patients to induce detergent-insoluble, pS129-positive immunofluorescence in neurons as a marker of αSyn pathology. Furthermore, we demonstrate that aggregates isolated from brain lysates from both PD and DLB patients also induce pS129 αSyn pathology, but only after prolonged incubation. We then used αSyn selective antibodies to immunodeplete the aggregate-inducing form from both PFF and synucleinopathy brain samples in order to understand which αSyn epitopes and species hold potential as therapeutic targets to prevent the pathological transmission of αSyn aggregates.

## Results

### Assay characterization using PFF

To determine if application of human αSyn PFF could induce a detergent-insoluble pS129 αSyn signal in primary rat neuronal cultures, hippocampal cells at 7 days in vitro (DIV) were treated with 150 ng/ml αSyn PFF generated from full-length recombinant hA53T-αSyn. This treatment led to a modest induction of pS129 αSyn immunofluorescence after 11 days in the absence of cellular toxicity (supplemental Figure S1). These results confirm previous reports that exogenous recombinant human αSyn PFF can induce the recruitment of endogenous rodent αSyn into insoluble pS129 αSyn aggregates (22, 23). In order to create a more sensitive assay system, primary rat hippocampal neurons were transduced with AAV-hA53T-αSyn at DIV 4 to overexpress hA53T-αSyn. Transduction with AAV-hA53T-αSyn alone did not induce accumulation of insoluble pS129 αSyn signal; however, application of 150 ng/ml PFF led to a robust induction of insoluble pS129 αSyn staining 11 days following treatment (Figure 1A-B). PFF-mediated induction in the AAV-hA53T-αSyn-transduced neurons was ~20-fold higher than induction in the neurons exposed to an empty AAV vector (Figure 1B). In addition, increasing the MOI of AAV-hA53T-αSyn led to an increase in the PFF-induced pS129 signal (supplemental Figure S2A) suggesting that the concentration of intra-cellular αSyn is a key parameter regulating induction by PFFs. Due to the report of cross-reactivity of the 81A anti-αSyn pS129 antibody with phosphorylated neurofilament light chain (NfL) (24), specificity of the 81A signal was confirmed by demonstrating a comparable staining pattern of insoluble pS129 with a second anti-pS129 antibody, MJFR13, which does not cross-react with phosphorylated NfL (supplemental Figure S3) (25).

**Figure 1.**
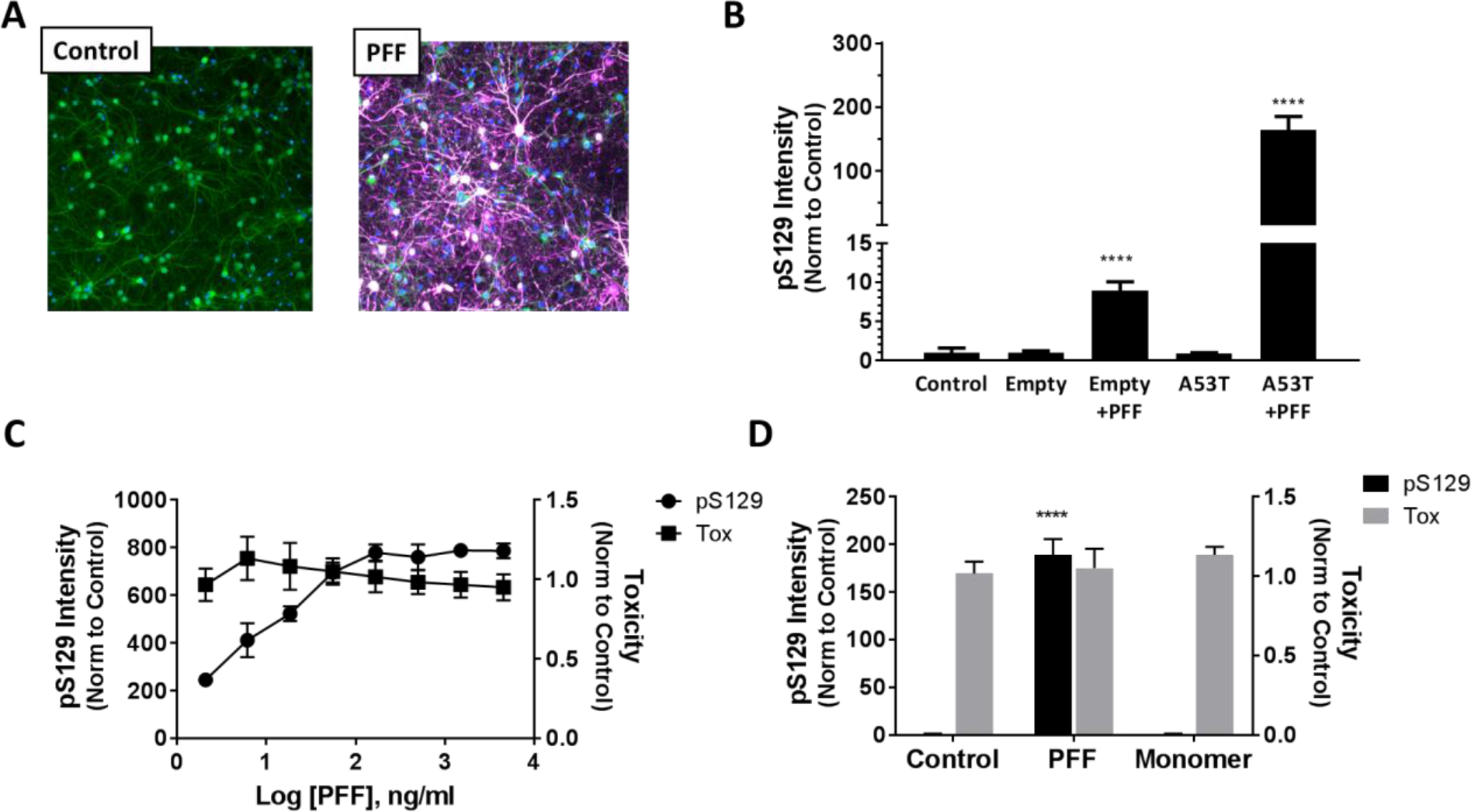
Characterization of pS129 αSyn induction with PFF. (A) Primary rat hippocampal neurons overexpressing hA53T αSyn were fixed in the presence of 1% Triton X-100 to extract soluble proteins 11 days (DIV 18) after treatment with 150 ng/ml PFF. Nuclei were stained with Hoechst (blue), dendrites with an anti-MAP2 antibody (green) and pS129 αSyn stained with antibody 81A (white/purple). Neurons treated with PFF exhibited robust pS129 αSyn staining compared to untreated controls. (B) The relative level of pS129 αSyn intensity normalized to untreated controls. Neurons were transduced with either an empty AAV (Empty) or A53T αSyn AAV (A53T) and then left untreated or treated with 150 ng/ml PFF for 11 days. Treatment with PFF resulted in a significant increase in pS129 signal in both empty AAV and hA53T αSyn AAV transduced neurons compared to control. Induction was ~ 20-fold higher in the hA53T αSyn AAV transduced neurons as compared to empty AAV transduced neurons. Data represents average ± SD from n=6 replicates. Statistical analysis based on one-way ANOVA with Dunnett’s post hoc test. (**** p < 0.0001). (C) αSyn PFF concentration response curve generated 11d post treatment in neurons transduced with hA53T αSyn AAV. A concentration-dependent increase in pS129 signal observed in the absence of cell toxicity. Data are normalized to untreated controls and represents average ± SD from n=6 replicate wells. (D) Neurons transduced with hA53T αSyn AAV were treated for 11 days with either 450 ng/ml PFF or 4500 ng/ml monomer and the induction of pS129 or the effect on cell health measured. Significant induction of pS129 signal observed with PFF but no induction with αSyn monomer. Data are normalized to untreated controls and represents average ± SD from n=6 replicates. Statistical analysis based on one-way ANOVA with Dunnett’s post hoc test. (**** p < 0.0001).

PFF-mediated induction of insoluble pS129 in AAV-hA53T-αSyn-transduced hippocampal neurons was concentration-dependent and did not cause significant toxicity (Figure 1C). In contrast, a pS129 αSyn signal could not be elicited with 4500 ng/ml of hA53T-αSyn monomer (Figure 1D) or with PFF of tau protein (data not shown). Furthermore, PFF-dependent increases in pS129 αSyn were inversely correlated with the level of branch points in MAP2-positive neurites (supplemental Figure S2B). These data are consistent with a previous publication reporting that overexpression of hA53T-αSyn in rat primary midbrain neurons results in altered neurite morphology, including decreased neurite branching (26). Our results demonstrate that detergent insoluble pS129 αSyn pathology can be induced with hA53T-αSyn PFF in primary rat hippocampal neurons overexpressing hA53T-αSyn and is correlated with impaired neurite morphology.

### Assay characterization using human brain lysates

Having developed a highly sensitive reporter assay, we next sought to determine if lysates generated from brain samples of patients diagnosed with various synucleinopathies could induce pS129 αSyn pathology. PBS-soluble lysates were generated using frontal cortex tissue samples from MSA, PD, DLB+AD patients and normal controls (demographic and neuropathological information are included in supplemental Table S1 and S2, respectively). AAV-hA53T-αSyn-transduced hippocampal neurons were treated with lysates and the induction of insoluble pS129 αSyn measured. MSA lysates induced a significant increase in pS129 αSyn signal 11 days post treatment in contrast to brain lysates from PD, DLB+AD and normal controls (Figure 2A-B, supplemental Table S3). Induction was robust (~100-fold of untreated control) and significant toxicity was not observed. A total of 9 different MSA brain lysates were then tested at different concentrations (Figure 2C, supplemental Table S4). Statistically significant induction was observed with 5 of the 9 MSA lysates tested at a 1:300 dilution, with the highest measured intensity (MSA #1, MSA #7) resulting in ~50% of the pS129 signal induced by a 150 ng/ml PFF treatment (Figure 2C, supplemental Table S4). Variability in the absolute degree of induction for MSA #1 and MSA #3 (compare Figure 2B and 2C) was observed and likely reflects inter-batch differences in the primary neuron cultures. Nevertheless, the rank order of the ability and degree to which individual MSA brain samples induced was consistent across several primary neuron preparations. MSA lysate induction was concentration dependent and lysates did not produce toxicity with the exception of the highest concentration tested (1:100; data not shown). These results are in agreement with a previous reports demonstrating that only lysates from MSA patients and not PD patients induce αSyn aggregation in HEK cells expressing αSyn A53T-YFP, and that different MSA brain lysates produce different magnitudes of induction (10, 27).

**Figure 2.**
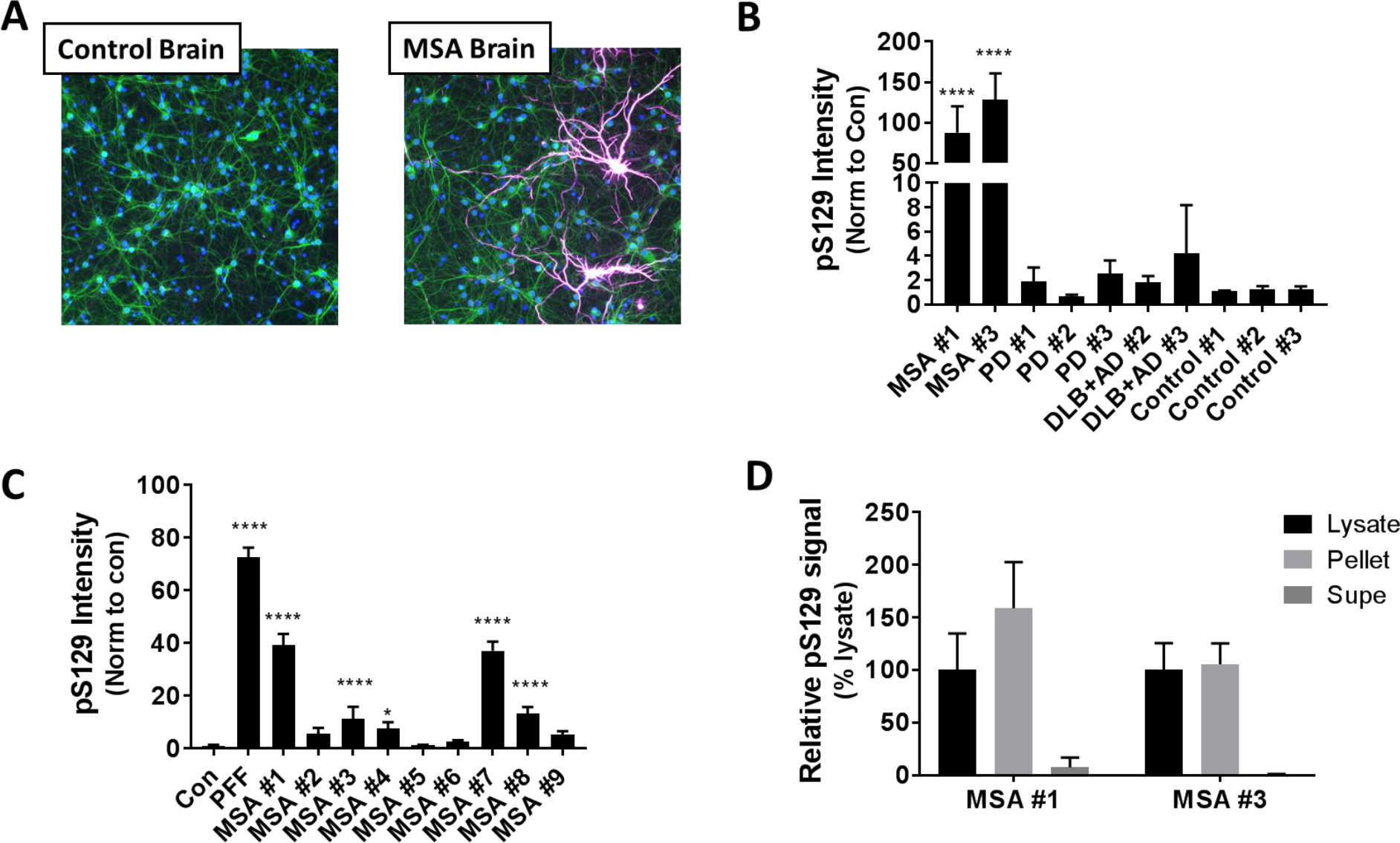
Characterization of pS129 αSyn induction with brain lysates. (A) Primary rat hippocampal neurons overexpressing hA53T αSyn were fixed in the presence of 1% Triton X-100 to extract soluble proteins and stained with Hoechst (blue), an anti-MAP2 antibody (green) and the anti-pS129 αSyn antibody 81A (purple) 11 days (DIV 18) after treatment with a 1:300 dilution of lysates from control brain and MSA brain tissue. (B) Quantification of pS129 αSyn intensity normalized to untreated control wells shows robust and statistically significant elevated levels of pS129 αSyn induction for neurons treated with MSA brain tissue lysates as compared to either control, PD or DLB+AD brain lysates. Data represent average ± SD from n= 6-12 replicates for untreated controls and 4-6 replicates for brain lysates. <5% of treated wells were excluded from analysis due to toxicity (toxicity index <0.6). Statistical analysis based on one-way ANOVA with Dunnett’s post hoc test. (**** p < 0.0001). (C) Brain tissue lysates from 9 different MSA patients showing varying amounts of pS129 αSyn induction 11days post treatment normalized to the untreated control. Data represent average ± SD from n=6 replicates. All wells exhibited acceptable viability scores (toxicity index > 0.6). Statistical analysis based on comparison to untreated controls using one-way ANOVA with Dunnett’s post hoc test. (*p<0.5; **** p < 0.0001). (D) MSA brain tissue lysates were fractionated by high-speed centrifugation and the resulting pellet and supernatant (Supe) fractions measured for pS129 induction. Induction is expressed relative to induction with the starting lysate. Data represent average ± SD from n=6 replicates.

The magnitude of induction by the MSA lysates did not correlate with αSyn levels (supplemental Figure S4A) suggesting that the inducing species may be a minor component of the overall αSyn in the extract. In addition, induction did not correlate with the extent of glial cytoplasmic inclusions (GCIs) observed in tissue sections, suggesting that there may be multiple variables affecting the level of the PBS-soluble inducing species (supplemental Figure S4B).

High-speed centrifugation has been used to partially-purify the pathology-inducing species of tau from brain lysates (28) and a similar approach was used to further characterize the inducing species from the MSA samples. PBS-soluble MSA brain lysates were subjected to high-speed centrifugation (100,000 x g, 30 minutes) and the resulting supernatant and pellet fractions isolated and tested for induction in transduced neurons. As shown in Figure 2D, inducing activity was completely recovered in the re-suspended pellets but not the supernatant fractions from the MSA lysates. This finding indicates that the inducing species can be enriched by high speed centrifugation and is likely composed of high molecular weight aggregates.

### Longer incubation times with PD and DLB+AD brain lysates lead to pS129 αSyn induction

Both the data presented here, as well as recently published data, suggest that MSA and PD may contain different strains of toxic αSyn species due to their differential ability to induce pathology in various biological systems (10, 27). Because of these potential strain differences, we hypothesized that DLB+AD or PD brain lysates may require longer incubation times for induction of pS129 αSyn pathology in our in vitro system. As with the MSA samples, high-speed centrifugation was used to enrich the presumed αSyn aggregates from control, PD and DLB+AD lysates. Levels of αSyn in starting brain lysates and isolated supernatant and pellet fractions were measured by ELISA (Figure 3, supplemental Table S5). As shown in Figure 3, similar levels of αSyn were detected across the starting lysates and the pellet fractions regardless of clinical diagnosis. On average ~3% of the starting αSyn ELISA signal was recovered in the pellet fraction while ~32% was recovered in the supernatant fraction (supplemental Table S5). The reasons for the modest recovery of total αSyn starting material (~35%) following centrifugation are not clear. These results indicate that high molecular weight species of αSyn, presumably aggregates, are present in all of the brain lysates, including control lysates. Furthermore these high molecular aggregates represent a minor fraction (~3%) of the total αSyn present in the starting lysate. Re-suspended pellets were then tested for inducing activity in AAV-hA53T-αSyn-transduced hippocampal neurons. Cells were treated for up to 32 days, 3 weeks longer than previous experiments. As expected, significant induction of pS129 was observed with MSA pellet fractions following 11 days of treatment (Figure 4, supplemental Table S6) and the induction was significant at all time points. The ~2-fold higher induction at 11 days compared to the later time points likely reflects inter-plate differences in background normalization. Interestingly, significant induction was also observed with the DLB+AD and PD pellet fractions but only after 25 days of incubation (Figure 4, supplemental Table S6). In contrast, pellet samples from control brain lysates failed to induce pS129 αSyn, regardless of duration of exposure (Figure 4, supplemental Table S6). As the total αSyn levels were similar across all of the pellet fractions (Figure 3), the observed differences in induction are likely related to differences in the molecular nature and/or conformation of the high molecular weight species enriched by high speed centrifugation.

**Figure 3.**
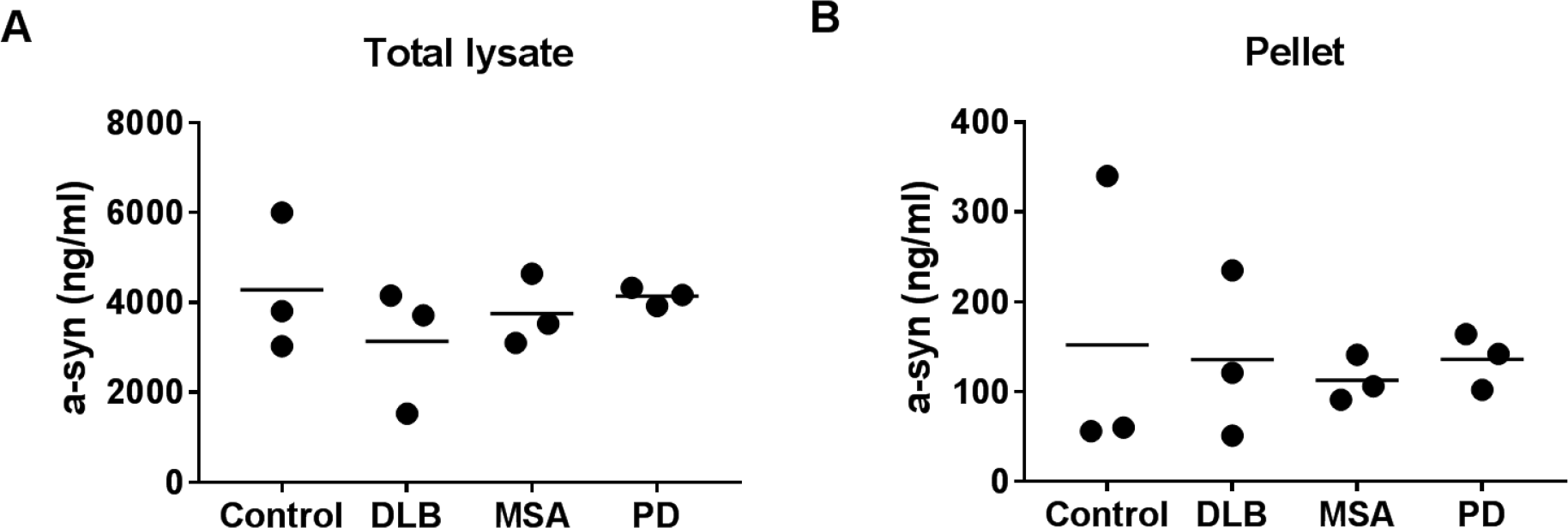
αSyn levels in brain lysates. αSyn levels were measured by ELISA in (A) total brain lysates and (B) aggregates isolated from brain lysates following high-speed centrifugation (Pellet). αSyn levels are expressed relative to a αSyn monomer standard. Each data point represents individual control, DLB, MSA and PD samples. Group averages are indicated. No significant differences between groups were observed based on one-way ANOVA.

**Figure 4.**
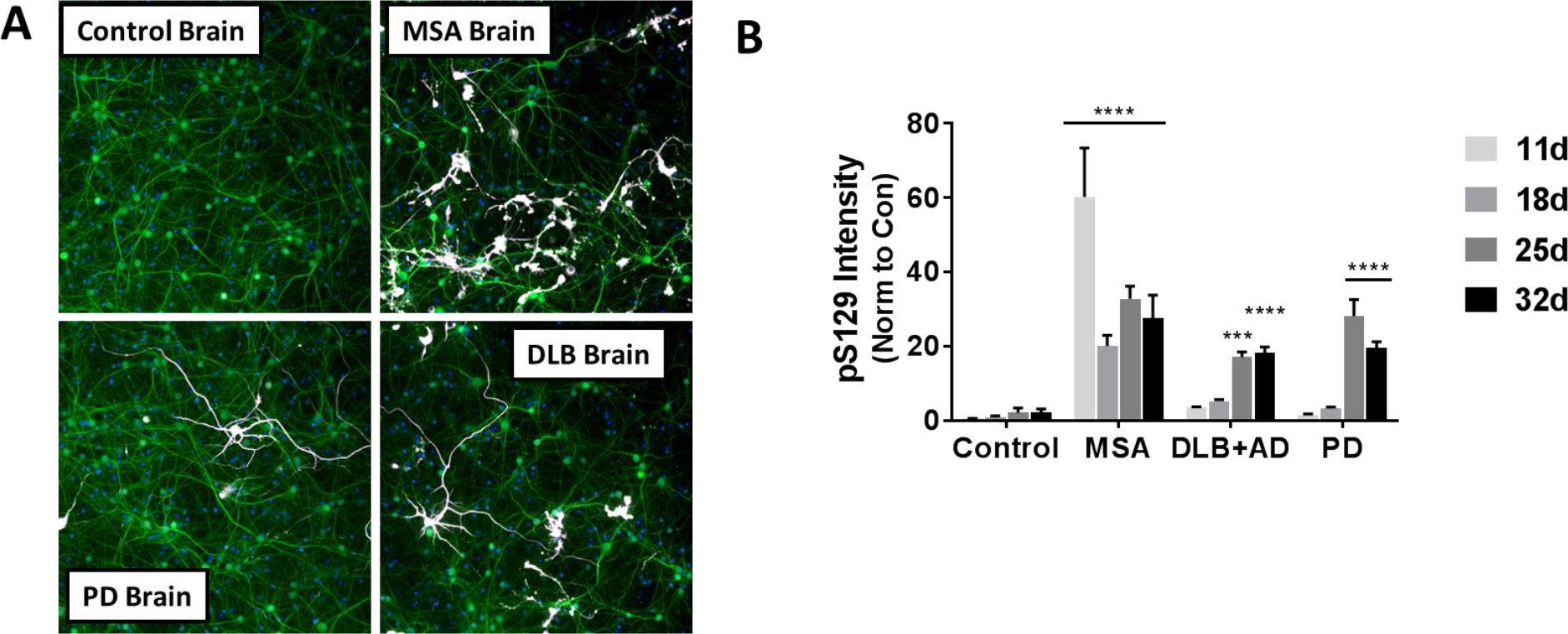
Induction of pS129 αSyn signal with pellet fractions from brain lysates. (A) Primary rat hippocampal neurons overexpressing hA53T αSyn were fixed in the presence of 1% Triton X-100 to extract soluble proteins and stained with Hoechst (blue), an anti-MAP2 antibody (green) and the anti-pS129 αSyn antibody 81A (white) 25-days (DIV 32) after treatment with re-suspended brain lysate pellet fractions from control, MSA, PD, DLB cases. (B) Quantification of pS129 αSyn intensity normalized to untreated controls following 11, 18, 25 or 32 days of treatment. Significant induction of pS129 αSyn can be seen for all three synucleinopathy brain tissue samples. No induction of pS129 αSyn signal was detected with control brain tissue pellets at any timepoint. Data represent average ± SD from 3 different brain lysate pellets for each group. Individual brain pellet data are included in supplemental Table S6. Statistical analysis based on comparison to the control group using two-way ANOVA with Dunnett’s post hoc test. (*** p < 0.001; **** p < 0.0001).

### Binding of αSyn antibodies to αSyn monomers and PFFs

To further investigate the molecular nature of the induction-competent species found in αSyn PFF and MSA brain lysates, a set of ten commercially available αSyn antibodies were selected for immunodepletion studies. Antibodies were selected to target different epitopes across the αSyn sequence (Figure 5, supplemental Table S7). A competitive binding assay was used to characterize antibody affinity to αSyn monomers and PFF and example binding curves for MJFR1 and MJFR14642 are shown in Figure 5. A summary of relative affinity IC_50_ values is included in Table 1. As shown in Figure 5B, MJFR1 exhibited comparable binding to both αSyn monomer and αSyn PFF with IC_50_ values of 8.0 ng/ml and 29.2 ng/ml, respectively, and a monomer to PFF binding ratio of 0.27 (Table 1). In contrast, MJFR14642 exhibited 50-fold higher affinity for PFF and over 400-fold weaker affinity for monomer compared to MJFR1 (Figure 5C, Table 1) demonstrating that MJFR14642 is more selective for the oligomeric form. Indeed, MJFR14642 exhibited the greatest degree of selectivity for PFF (monomer to PFF binding ratio of 7173) of all the antibodies tested (Table 1). Only 4D6 and LB509, two other C-terminal antibodies, produced monomer to PFF binding ratios greater than 10 (26 and 39, respectively; Table 1).

**Figure 5.**
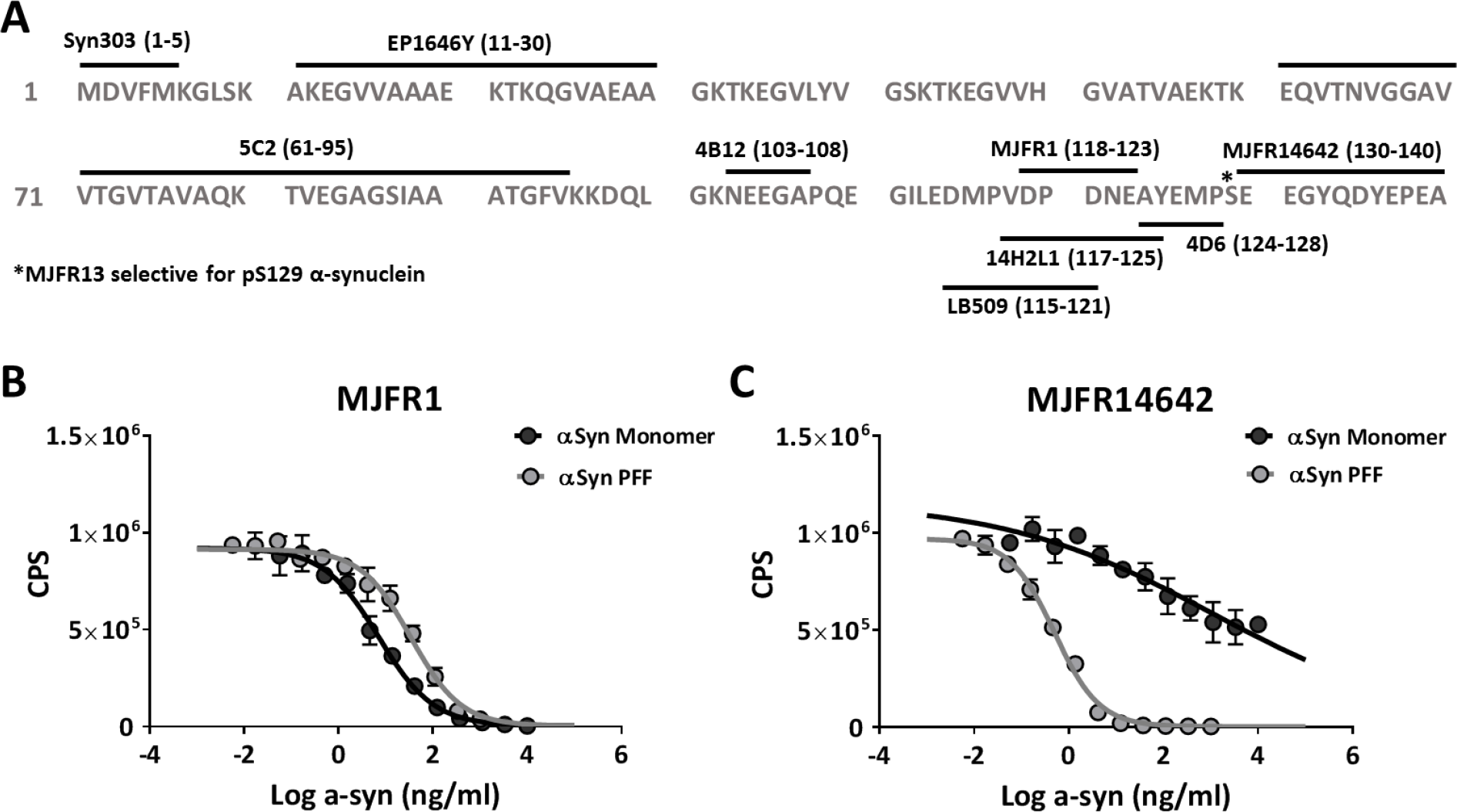
Binding characterization of αSyn antibodies. (A) Amino acid sequence for αSyn showing the putative binding regions for the commercial αSyn antibodies tested. *S129 residue. (B) Example curves for MJFR1 binding to increasing concentrations of either αSyn monomer (black) or αSyn PFF (gray). Data represent average ± SD from duplicate wells. Summary IC_50_ values included in Table 1. (C) Example curves for MJFR14642 binding to increasing concentrations of either αSyn monomer (black) or αSyn PFF (gray). Data represents average ± SD from duplicate wells. Summary IC_50_ values included in Table 1.

**Table 1.** Antibody binding IC_50_ values

| Antibody | Monomer Binding (ng/ml) |  |  | PFF Binding (ng/ml) |  |  | Mono/PFF<br>IC<sub>50</sub> Ratio |
| --- | --- | --- | --- | --- | --- | --- | --- |
|  | AVE | SD | n | AVE | SD | n |  |
| MJFR 14642 | 3658 | 1981 | 5 | 0.51 | 0.10 | 5 | 7173 |
| MJFR1 | 8.0 | 1.3 | 2 | 29.2 | 7.0 | 2 | 0.27 |
| 4D6 | 54 | 18 | 2 | 2.1 | 0.2 | 2 | 26 |
| 4B12 | 5.6 | 1.2 | 2 | 23.3 | 3.2 | 2 | 0.24 |
| Syn303 | 51 | 9 | 2 | 109.3 | 6.6 | 2 | 0.47 |
| LB509 | 62 | 24 | 2 | 1.6 | 0.1 | 2 | 39 |
| EP1646Y | 17 | 0.4 | 2 | 52.8 | 9.9 | 2 | 0.32 |
| 5C2 | >10000 |  | 2 | >1000 |  | 2 | NA |
| 14H2 | 21 | 5 | 2 | 10 | 3 | 2 | 2.1 |

Average IC<sub>50</sub> for inhibition of binding of antibodies to PFF-coated plates by αSyn monomers and αSyn PFF. Antibodies were preincubated in solution with a titration of either αSyn monomer or PFF and then the amount of free antibody determined and IC<sub>50</sub> calculated. Data represent the average ± SD from 2-5 experiments. The monomer to PFF IC<sub>50</sub> ratios are also shown.

### Immunodepletion of PFF and MSA-induced pS129 αSyn pathology with αSyn antibodies

The ability of each antibody to immunodeplete the αSyn inducing species from either αSyn PFF or a MSA brain lysate sample was then evaluated. Antibodies were tested at a concentration of 50 nM. As we were unable to demonstrate binding of antibody 5C2 to either monomer or PFF up to 15000 ng/ml (Table 1), this antibody, along rabbit IgG, were included as negative controls. PFF αSyn or MSA brain lysate sample #7 were incubated overnight with each antibody, the antibody complexes removed and the immunodepleted samples tested for induction of pS129 αSyn in AAV-hA53T-αSyn-transduced hippocampal neurons after 11 days of treatment (Figure 6). The greatest depletion of the inducing species from PFF was observed with the C-terminal antibodies LB509, 14H2, MJFR1 and MJFR14642 with reductions of >90%. In contrast, the N-terminal antibodies Syn303 and EP1646Y exhibited limited ability to immunodeplete the PFF inducing species, producing reductions of ~30% and ~50%, respectively. This result is consistent with the lower activity of Syn303 compared to the C-terminal antibody Syn211 in inhibiting PFF-mediated induction of pathology in mouse neurons as previously reported (23). The pS129-selective antibody MJFR13 was unable to immunodeplete the inducing species from PFF, consistent with the lack of pS129 αSyn in the PFF samples.

**Figure 6.**
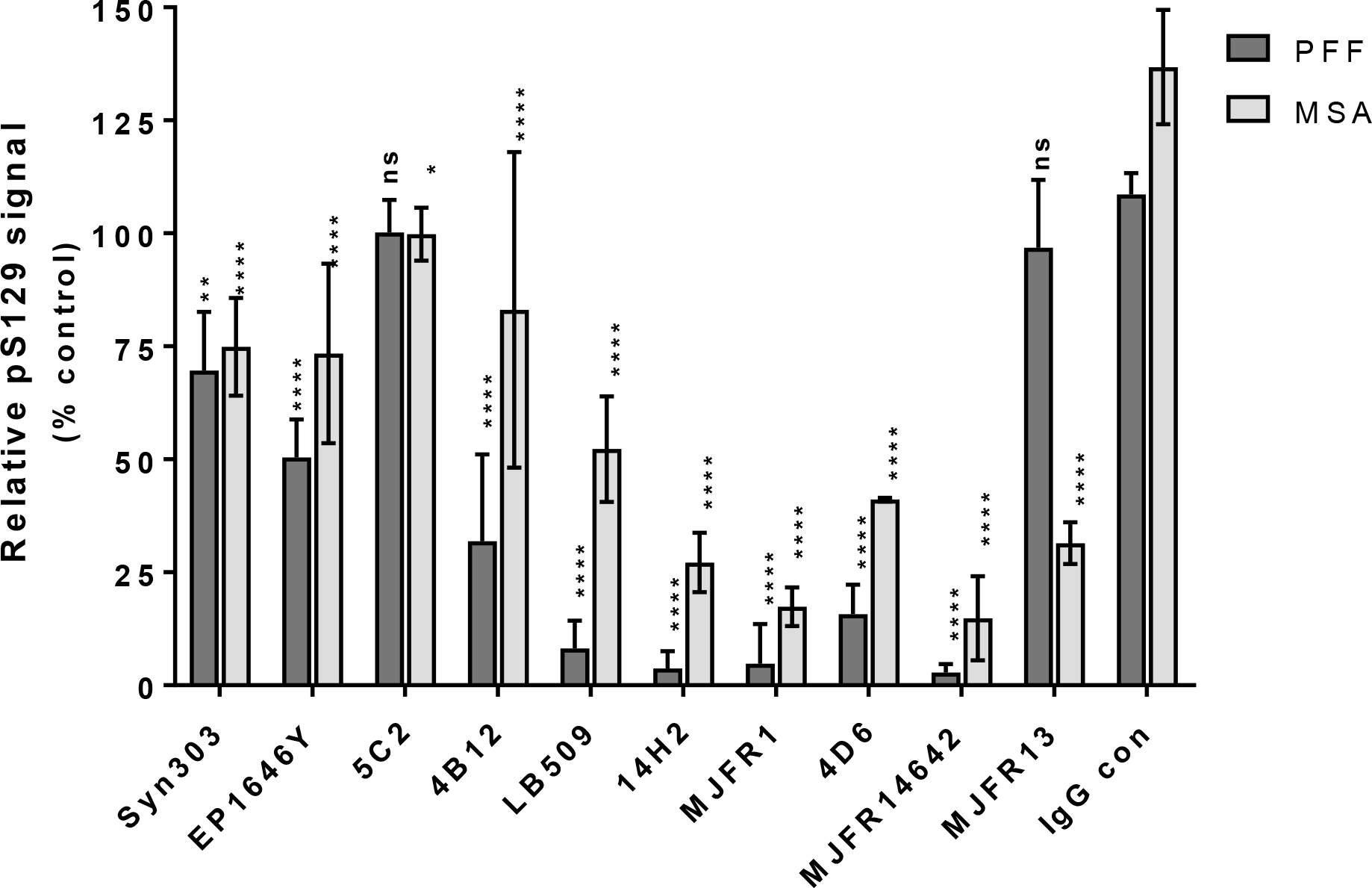
Immunodepletion of inducing activity from PFF and a MSA brain lysate with αSyn antibodies. PFF (15 ng/ml) or MSA brain lysate #7 were immunodepleted with αSyn antibodies at a concentration of 50 nM. Depleted PFF or brain lysate samples were then tested for induction of pS129 αSyn in primary rat hippocampal neurons overexpressing hA53T αSyn following 11 days of treatment. Data are normalized to the undepleted PFF or undepleted MSA lysate positive controls and represents the average ± SD from 2-6 experiments. Statistical analysis based on comparison to the IgG control group using two-way ANOVA with Dunnett’s post hoc test. (* p < 0.05; ** p < 0.01; **** p < 0.0001; ns = not significant).

MJFR1 and MJFR14642 were the most active antibodies for immunodepleting the induction-competent species from the MSA brain lysate with reductions of >80% observed (Figure 6). Interestingly, LB509, another C-terminal antibody, exhibited limited ability to deplete the inducing species from the MSA lysate (~50%) even though this antibody depleted >90% of the inducing species from PFF. The lower than expected activity of LB509 for depleting the inducing species from MSA could reflect potential differences in the conformation and/or modifications of the MSA species compared to PFF. Similar to the results with PFF, both N-terminal antibodies were less effective for depleting the inducing species from MSA. In contrast to the results with PFF, the pS129 antibody MJFR13 immunodepleted the MSA αSyn inducing species by ~70%, indicating the presence of pS129 αSyn. This observation is intriguing considering that phosphorylated oligomeric αSyn has been shown to be significantly elevated in the CSF of MSA patients as compared to other synucleinopathies (29).

Since both MJFR1 and MJFR14642 were highly effective for depleting the inducing species from PFF and the MSA lysates, these antibodies were selected for concentration response curve (CRC) analysis (Figure 7). Both antibodies exhibited similar maximal efficacy and potency in depleting the PFF inducing species (Figure 7A; IC_50_ values derived from several experiments are included in Table 2); IC_50_ values of 0.036 nM and 0.079 nM were derived for MJFR1 and MJFR14642, respectively. To determine if differences in oligomer selectivity influenced the ability of each antibody to deplete the inducing species in the presence of monomer, PFF immunodepletion reactions with either 1 nM MJFR1 or 1 nM MJFR14642 were spiked with increasing concentrations of αSyn monomer and the depleted samples tested for inducing activity (Figure 7C). In the absence of spiked monomer, both antibodies completely depleted the inducing species from PFF as expected. Addition of αSyn monomer effectively blocked the ability of MJFR1 to deplete the inducing species in PFF in a dose-dependent manner (Figure 7C). In contrast, MJFR14642 was ~100-fold less sensitive to competition with αSyn monomer (Figure 7C). Taken together, these results confirm the superior selectively of MJFR14642 and suggest that this property may translate into more potent activity against the induction-competent species in biological fluids that contain competing levels of αSyn monomer (e.g. CSF and ISF).

**Figure 7.**
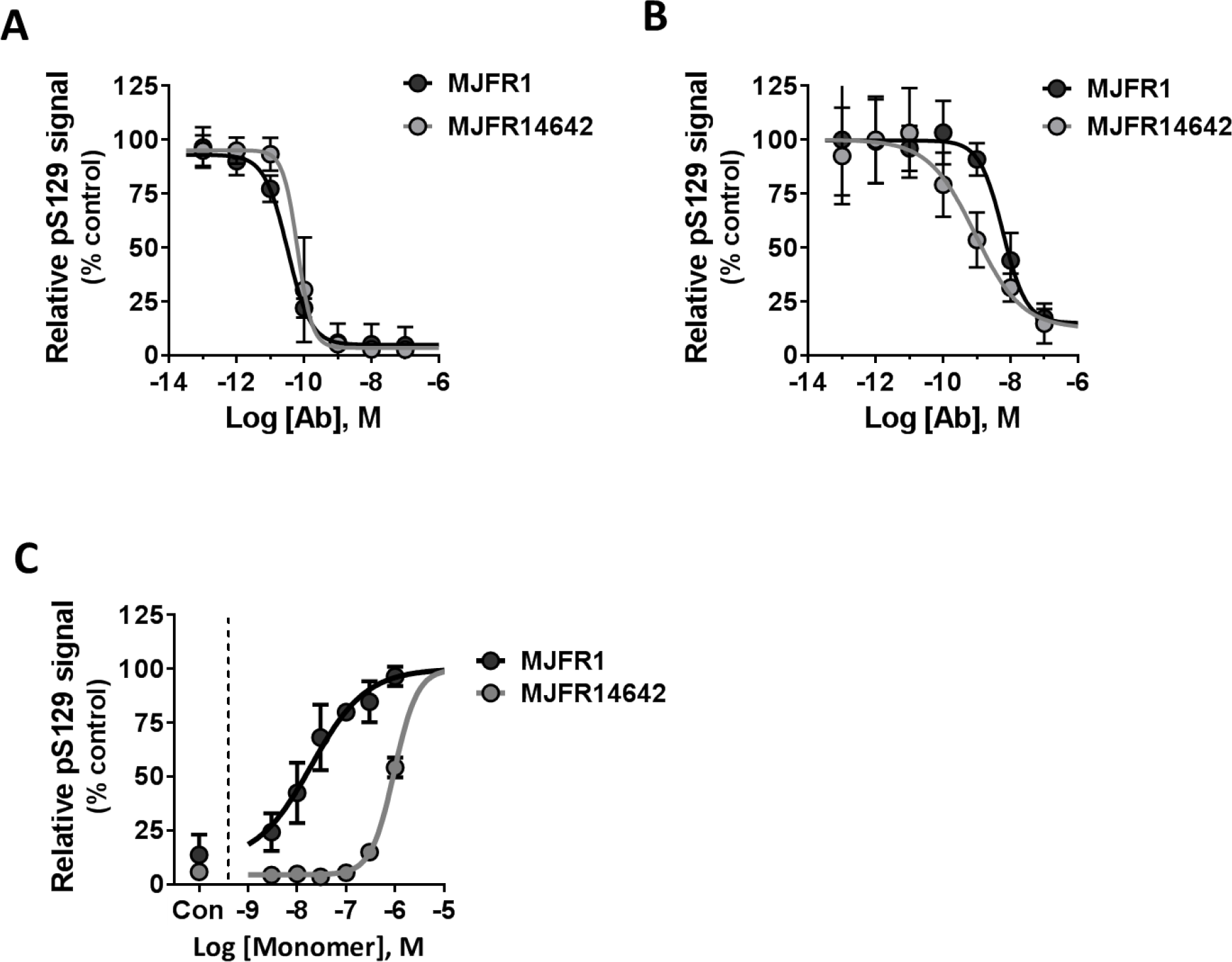
Immunodepletion of PFF and a MSA brain tissue sample with antibodies MJFR1 and MJFR14642. Samples were preincubated with a titration of either MJFR1 or MJFR14642, the antibody-complexes removed and the immundepleted samples evaluated for induction of pS129 αSyn pathology in primary rat hippocampal neurons overexpressing hA53T αSyn following 11 days of treatment. (A) Example immunodepletion curves for MJFR1 (black) and MJFR14642 (gray) for removing the inducing αSyn species from 15 ng/ml PFF. Data was normalized to an undepleted PFF positive control. Cumulative curves were generated by combining data from multiple experiments. Data represent average ± SD from n=4 experiments. Summary IC_50_ values are included in Table 2. (B) Example immunodepletion curves for MJFR1 (black) and MJFR14642 (gray) for removing the inducing αSyn species from MSA brain tissue lysates #7. Data were normalized to an undepleted MSA brain lysate as positive control. Cumulative curves were generated by combining data from multiple experiments. Data represent average ± SD from n=3-6 experiments. Summary IC_50_ values are included in Table 2. (C) Example immunodepletion curves for 1 nM MJFR1 (black) and 1 nM MJFR14642 (gray) for removing the inducing αSyn species from 15 ng/ml PFF samples in the presence of increasing concentrations of αSyn monomer. Data were normalized to an undepleted PFF positive control. Data represent average ± SD from n=3 replicates.

**Table 2.** Antibody immunodepletion IC_50_ values

| Sample | MJFR1 IC<sub>50</sub> (nM) |  |  | MJFR14642 IC<sub>50</sub> (nM) |  |  | R1/R14642<br>IC<sub>50</sub> ratio |
| --- | --- | --- | --- | --- | --- | --- | --- |
|  | AVE | SD | n | AVE | SD | n |  |
| PFF | 0.036 | 0.017 | 4 | 0.079 | 0.054 | 4 | 0.5 |
| MSA #7<br>lysate | 7.2 | 2.2 | 3 | 0.72 | 0.38 | 5 | 10.0 |
| MSA #7<br>pellet | 0.52 |  | 1 | 0.038 |  | 1 | 13.7 |
| PD #3<br>pellet | 0.29 |  | 1 | 0.021 |  | 1 | 13.8 |
| DLB+AD #2<br>pellet | 0.44 |  | 1 | 0.062 |  | 1 | 7.1 |

Average IC<sub>50</sub> for immunodepletion of the induction-competent species from PFF, MSA, PD and DLB+AD brain extracts. Samples were preincubated with a titration of either MJFR1 or MJFR14642, the antibody-complexes removed and the immunodepleted samples evaluated for induction of pS129 αSyn pathology in primary rat hippocampal neurons overexpressing hA53T αSyn following 11 days of treatment. In some cases, the inducing species was enriched by high-speed centrifugation (pellets) prior to immunodepletion. Data represent the average ± SD from 3-5 experiments or the IC<sub>50</sub> from a single experiment. The MJFR1 to MJFR14642 IC<sub>50</sub> ratios are also shown.

MJFR1 and MJFR14642 also depleted the inducing species from MSA lysate #7 in a concentration dependent manner (Figure 7B). Both antibodies were weaker for immunodepleting the inducing αSyn species from the MSA lysate compared to PFF (Table 2). Interestingly, the oligomer-selective antibody MJFR14642 was ~10-fold more potent in depleting the MSA inducing species compared to the non-selective MJFR1 antibody (Table 2). This ~10-fold difference in potency was also maintained when the MSA brain pellet was used to induce αSyn aggregates (13.7 fold, Table 2, N=1). This finding suggests that the MJFR14642 vs. MJFR1 potency difference is likely due to differences in conformation and/or epitope exposure of the induction-competent species, rather than competitive binding to other non-inducing αSyn found in the crude brain lysates (e.g., αSyn monomers).

Both αSyn antibodies MJFR1 and MJFR14642 were also able to immunodeplete the inducing species from pelleted PD and DLB brain lysate samples in a concentration dependent manner (Table 2). Interestingly, the αSyn oligomer-selective MJFR14642 exhibited 7.1-fold and 13.8-fold greater potency than MJFR1 for immunodepleting the inducing αSyn species from a DLB brain pellet and a PD brain pellet, respectively (Table 2). These data confirm that induction of pS129 αSyn by PD and DLB extract pellets is mediated by αSyn aggregates and further support the hypothesis that C-terminal, oligomer-selective αSyn antibodies are more specific for the physiologically-relevant transmissible forms of αSyn from synucleinopathy patients.

## Discussion

The hypothesis that the pathological initiation and progression of synucleinopathies is in part due to cell to cell transmission of toxic αSyn species in a prion-like fashion has gained support in recent years (8, 9). As such, the concept of an immunotherapy in which αSyn-selective antibodies bind to and prevent the spread of the pathological αSyn species responsible for templating toxic αSyn aggregates has become an attractive disease-modifying hypothesis and approach (30). Our data demonstrate that αSyn antibodies can bind to induction-competent αSyn species in synthetic αSyn PFFs and pathologically-relevant αSyn aggregates found in brain lysates from synucleinopathy patients, thereby preventing the induction of insoluble pS129 αSyn-positive aggregates in primary rat hippocampal cultures.

There are three main findings from the experiments described in this report. First, we demonstrate that frontal cortex lysates from synucleinopathy patients induce pathological pS129 αSyn in a rodent primary neuron system. While pS129 αSyn induction has been shown previously in rodent primary cultures using synthetic αSyn PFFs (22, 23), and brain tissue samples from MSA patients have been reported to induce αSyn aggregates in αSyn-expressing HEK cells (10, 27), this is the first report to demonstrate the ability of MSA brain tissue samples to induce insoluble pS129-positive aggregates of αSyn in primary neuronal cultures. Moreover, while our initial data demonstrating a lack of induction of pS129 αSyn following 11 days of treatment with either PD or DLB brain lysates agree with previous data showing the inability of PD brain tissue to induce αSyn aggregates in αSyn-expressing HEK cells (10, 27), we demonstrate for the first time that PD and DLB brain tissue samples can induce a pathological signal in neurons after prolonged incubation times. This result is especially intriguing given the clinical observation that MSA typically progresses much more rapidly than PD and DLB (31). This finding is consistent with the idea that different toxic αSyn strains are present contingent on the differing synucleinopathy clinical and pathologic states (32, 33). Furthermore, the current data support the hypothesis that different conformations and/or posttranslational modifications can confer varying degrees of αSyn strain toxicity or infectivity, which may in turn drive the rate of disease progression (10, 32, 34, 35). The potential differences between the discrete αSyn strains have yet to be clearly elucidated. There are reports of different amounts of SDS-soluble and insoluble αSyn species in PD, DLB and MSA (36, 37) but the significance of these findings is unclear and additional work is required to identify the specific conformational and/or posttranslational differences across toxic αSyn strains. Finally, while the strain hypothesis is an attractive explanation for the observed temporal differences in induction between MSA, PD and DLB, we cannot rule out the possibility that there is simply a greater concentration of the same strain of induction-competent αSyn in MSA relative to PD and DLB brain lysates.

A second finding is the superior potency and maximal efficacy of C-terminal antibodies in immunodepleting toxic αSyn species from diseased brain tissue samples. The potential for αSyn immunotherapy in synucleinopathies with C-terminal antibodies has been demonstrated in transgenic mouse models overexpressing αSyn (36–38), and immunotherapy with C-terminal antibodies has been proposed as a preferred strategy for targeting exposed regions of aggregated αSyn in order to inhibit fibrillization and membrane permeabilization (39). Here we report that C-terminal antibodies exhibit higher potency to immunodeplete induction-competent αSyn species present in MSA patient brain tissue compared with antibodies targeting the N-terminal region of αSyn. The C-terminal is the site of S129 phosphorylation and also contains a protease cleavage site at D119 that allows for the generation of a C-terminal truncated form of αSyn (40). The C-terminal truncated form of αSyn has been shown to be both a component of αSyn lesions (41, 42) and to enhance oligomerization and toxicity (41, 43–45). Furthermore, it has been shown that preventing C-terminal truncation with antibodies directed to the cleavage site is protective both in vitro and in vivo (46, 47). Of the 5 C-terminal antibodies tested here, 3 have epitopes that overlap this cleavage site (MJFR1, 14H2L1, LB509), while the other 2 are carboxy to this site (MJFR14642, 4D6). It is interesting to note that the most potent antibody for immunodepletion of the inducing αSyn species in MSA brain tissue, MJFR14642, binds to an epitope at the most distal region of αSyn and carboxy to the cleavage site. This observation suggests that while C-terminal truncated forms of αSyn may initiate the formation of and/or comprise a part of the inducing species in MSA brain tissue, an αSyn species with an intact C-terminal must be a major component of the induction-competent αSyn species found in the brains of MSA patients.

A third main finding is the increased potency of the highly oligomer-selective antibody MJFR14642, compared with the non-selective antibody MJFR1, for immunodepletion of the induction-competent αSyn species present in MSA, PD and DLB brain tissue samples. While both of these antibodies were equipotent and equally efficacious for immunodepleting the induction-competent αSyn species in PFF samples, MJFR14642 exhibited a ~10-fold greater potency for immunodepleting the induction-competent αSyn species in synucleinopathy brain samples. The reasons for the increased potency are unclear but may be related to a mixture of inducing species in the MSA samples for which the antibodies have varying affinity, or improved avidity of MJFR14642 for the inducing species. One limitation of this current study is that estimates of potency with the clinical samples are based on a limited number of samples and replicates. Nevertheless, these data support the notion of differences between inducing αSyn species in PFF and MSA brain tissue samples, as well as suggesting that αSyn oligomer-selective antibodies may be a preferred immunotherapy for synucleinopathies compared to αSyn antibodies that do not distinguish between conformations. Several oligomer-selective antibodies have been generated (48, 49), indicating the feasibility of generating antibodies targeting specific αSyn conformations. Moreover, due to the relatively high plasma and CSF concentrations of soluble αSyn monomer compared with the oligomeric / aggregated form of αSyn (50–52), the binding of oligomer-selective antibodies to the toxic αSyn species would be substantially less hindered due to low affinity for the more prevalent monomeric species. Reduced binding to monomeric αSyn represents at least two advantages for oligomer-selective antibodies. First, the probability of interference by endogenous monomeric αSyn function is reduced and second, clinical doses of oligomer-selective antibodies would be predicted to be lower than non-selective antibodies. This point is critical in terms of low brain to plasma exposures seen with antibody treatments, with approximately 0.1-0.3% brain penetration typically reported (53). High concentrations of free antibody will be required to achieve relevant concentrations in brain parenchyma. Therefore, αSyn antibodies that are selective for αSyn oligomers and bind less αSyn monomer in plasma and the interstitial space will result in a greater fraction of free antibody available to access the CNS, reach therapeutic levels in the brain and neutralize oligomeric species in the interstitial fluid.

Finally, the in vitro system used in this study represents a robust assay for detecting pathological induction with relatively low amounts of the toxic αSyn species. However, the limitations of the system include that it is a rat rather than human preparation, that overexpression of mutant human αSyn is required to produce a pS129 αSyn signal with sufficient signal-to-noise to enable antibody potency estimates and that incubation with diseased brain parenchyma lysates rather than interstitial fluid are used as the source of inducer. Nevertheless, the assay system also includes unique strengths. It is a relatively sensitive assay conducive for identifying potent αSyn antibodies that recognize pathologically-relevant αSyn aggregates, it could serve as a potential biomarker assay if optimized for CSF samples in terms of both patient selection and target engagement studies for clinical trials, and it holds promise to further explore the process of αSyn aggregate uptake and seeding mechanisms (54). Finally, it would be of interest to investigate potential functional consequences of this robust pS129 αSyn signal in future studies to determine if changes in neuronal activity either precede or follow these phosphorylation events. Indeed, alterations in network connectivity have been previously reported in PFF-treated wildtype mouse hippocampal neurons as early as 4 days post treatment (22).

In conclusion, we have demonstrated discrete pS129 αSyn pathology in rodent primary neurons following exposure to MSA, PD and DLB brain tissue, and that antibodies targeting the C-terminal of αSyn are most effective at immunodepleting these potentially distinct transmissible forms of αSyn. We propose that C-terminal antibodies selective for oligomeric forms over monomeric forms of αSyn represent a desirable immunotherapy for synucleinopathies due to immunodepletion of pathologically-relevant forms of transmissible αSyn species found in the diseased brain of synucleinopathy patients.

## Materials and Methods

Chemicals, reagents and kits were purchased from ThermoFisher (Waltham, MA) unless indicated otherwise.

### Preparation of α-synuclein pre-formed fibrils (PFFs)

Recombinant human αSyn (rPeptide; Bogart, GA) and human αSyn containing the A53T mutation (rPeptide; Bogart, GA) were reconstituted with H_2_O to a final concentration of 1 mg/ml in 20 mM Tris/HCl, 100 mM NaCl, pH 7.4. Samples were incubated in 2 ml eppendorf tubes (~1 ml/vial) at 37°C with constant shaking for 4 days and then centrifuged at 100,000 x g at room temperature (RT) for 20 minutes (min). The pellets were re-suspended with PBS for a final PFF concentration of 1 mg/ml (PFF concentrations expressed as monomer equivalent). PFF aliquots were frozen in liquid nitrogen and stored at −20°C. Fibrillization of αSyn was monitored by a Thioflavin T binding assay (supplemental Figure S5). Briefly, samples were diluted to 0.5 mg/ml in PBS and added to an equal volume of 25 μM Thioflavin-T. Samples were then measured using an Envision multilabel plate reader (PerkinElmer; Waltham, MA) with the excitation and emission wavelengths set at 485 nm and 535 nm, respectively. Total protein levels were assessed with a Micro BCA Protein Assay Kit.

### Preparation of human brain lysates

Frontal cortex brain tissue samples from normal controls and Multiple System Atrophy (MSA) and Dementia with Lewy Bodies plus Alzheimer’s disease (DLB+AD) cases were obtained from the Banner Sun Health Research Institute (Sun City, AZ). Frontal cortex brain tissue samples from Parkinson’s disease (PD) cases and normal controls were obtained from John Q. Trojanowski (University of Pennsylvania, Philadelphia, PA). Demographic information and neuropathology for each case are included in supplemental Tables S1 and S2, respectively. Brain samples were sonicated in filtered PBS (1 ml PBS/100 mg weight tissue) with a KONTES Micro Ultrasonic Cell Disrupter for 2 × 10 seconds (sec). Samples were placed in 2 ml eppendorf tubes and the tubes were kept on wet ice during sonication. Brain lysates were centrifuged at 3,000 x g, 4°C for 5 min to remove particulate material. The supernatant (PBS-soluble) was isolated, divided into aliquots and frozen in liquid N_2_ and stored at −80°C.

### High-speed centrifugation isolation of human brain lysates

Sedimentation of high molecular weight αSyn aggregates was adapted from (28). PBS-soluble brain homogenates previously prepared at 100 mg/ml in PBS were diluted 3-fold to 33.3 mg/ml in ice cold PBS followed by centrifugation at 100,000 x g for 30 min at 4°C. The supernatant (Supe) was removed and saved for analysis. The pellet was re-suspended in ice cold PBS in the same volume as the starting sample.

### Total and oligomer α-synuclein binding assay

Oligomer and monomer binding assays were performed essentially as described (49). Briefly, 96 well plates (Corning CoStar) were coated with 100 μl of 1 μg/ml αSyn WT PFF in PBS at RT for 2 hours (hr) or 4°C overnight (similar results were obtained with both). The plates were then washed 3 times with ~300 μl of wash buffer (0.05%Tween in PBS) and blocked with 150 μl of 3% BSA (Fraction V, protease-free, Roche Diagnostics Corporation; Indianapolis, IN) in PBS at room temperature for 2 hr (or 4°C overnight). 3-fold serial dilutions of PFF were prepared (final concentration range 1 μg/ml to 6 pg/ml, monomer equivalent) and αSyn WT monomer (final concentration range 10 μg/ml to 60 pg/ml) for 12 concentrations in sample buffer (0.1% BSA/0.05% Tween/PBS, 2 tablets of cOmplete Protease Inhibitor Cocktail (Roche Diagnostics Corporation; Indianapolis, IN) in 50ml buffer). An equal volume of 2-fold concentrated αSyn PFF or monomer was mixed with 2-fold concentrated antibodies in low binding plates (Becton Dickinson; Franklin Lakes, NJ) and incubated at room temperature for 2 hr. 100 μl of mixtures of antibody and PFF/monomer were loaded into the PFF coated plate and incubated at room temperature for 10 min, then washed 3 times. 100 μl of donkey anti-human IgG (Jackson ImmunoResearch; West Grove, PA), with 50% glycerol 1:1000 diluted in PBSTB (1% BSA/0.2% Tween/PBS) were then added and incubated at RT for 1 hr. Plates were washed 3 times for 5~10 min per wash. 100 μl of AP substrate (Tropix CDP Star Ready-to-Use with Sapphire II, Applied Biosystems; Waltham, MA) were then added and the reaction was developed at RT for 30 min. Luminescence counts were read with EnVision. PFF was sonicated 15-times at 1 sec/pulse before coating or mixing with antibodies. The plates were maintained with constant shaking during the assay.

### Total αSyn ELISA

Corning High Binding 96-well ELISA plates were coated with MJFR1 (Abcam, Cambridge, United Kingdom) (0.25 μg/ml) in Tris-buffered saline (TBS) (50 μl/well) for 1 hr at 37°C. Plates were then washed 4-times with 300 μl/well of TBS with 0.05% tween-20 (TBST) and then blocked for at least 2 hrs with 3% BSA/TBS at RT with shaking. The plates were washed as above followed by addition of the standard curve of synuclein peptide prepared in sample diluent (1% BSA/TBST). The highest standard concentration was 1000 pg/ml. Test samples were diluted in sample diluent, added at 50 μl/well to the ELISA plates and the plates incubated overnight at 4°C. After the overnight incubation, the secondary antibody (Clone-42, BD Biosciences, San Jose, CA), previously conjugated to alkaline phosphatase (AP) using EZ-link AP conjugation kit (Novus Biologicals, LLC, Littleton, CO), was added to the ELISA plates at 50 μl/well without washing and incubated for 1 hr at RT. Plates were then washed 4-times with 300 μl/well TBST and prior to addition of 100μl/well Tropix CDP star ready substrate. Signal was measured using a TopCount (PerkinElmer, Waltham, MA). Data analysis was performed using Graphpad Prism (Graphpad Software, Inc. La Jolla, CA).

### Primary cell culture isolation

Primary rat hippocampal neuronal cultures were prepared weekly from ~7 litters at embryonic day 19 (E19) using the Papain Dissociation System, according to manufacturer’s instructions (Worthington Biochemical; Lakewood, NJ). Rat hippocampal cells were plated on poly-D lysine coated 96 well imaging plates (BD Biosciences, San Jose, CA) (~16 plates per week) at 30,000/well in neuronal culture medium (neural basal medium (NBM) containing 0.5 mM GlutaMax and B-27 supplement (Invitrogen; Carlsbad, CA).

### Immunoprecipitation experiments

Previously prepared brain homogenate (100 mg/ml) was diluted 150-fold in neuronal culture medium. Protein A/G agarose beads were prewashed once with PBS + 0.05% tween-20, then 3-times with PBS and then blocked with PBS + 1% BSA for 2 hr at 4°C prior to use. Test αSyn antibodies were added to the diluted brain homogenate and samples incubated for 2 hr at 4°C with end-over-end rotation. Prewashed and preblocked Protein A/G agarose bead slurry was then added at a 1:10 dilution and samples incubated overnight at 4°C with end-over-end rotation. Samples were then centrifuged at 1500 x g for 2 min to pellet the beads. The depleted supernatant was then removed and used for treatment in the primary neuron induction assay.

### Immunofluorescence assay

On day *in vitro* (DIV) 4, rat hippocampal neurons were transduced with an adeno-associated viral vector, AAV1 containing the cDNA for human αSyn harboring the A53T mutation (GeneDetect; Sarasota, FL) at a multiplicity of infection (MOI) of 3,000 unless indicated otherwise. On DIV 7 cells were treated with test samples (6 wells per treatment). All treatments were added by half medium exchange with 2-fold concentrated samples. Each plate contained a negative control (no treatment condition) and positive controls including 150 ng/ml PFF or a non-depleted brain extract. On DIV 18 (11 days post treatment), cells were fixed and stained for either total αSyn (soluble and insoluble) or insoluble αSyn or pS129 αSyn. Neurons were fixed with a solution containing 4% paraformaldehyde, 4% sucrose and either 0.1% Triton X-100 (Sigma-Aldrich, Saint Louis, MO) to fix all αSyn species, or 1% Triton X-100 to fix only the insoluble forms. Fixative was added to the cells for 15 min. Following fixation, cells were washed 3-times with wash buffer containing PBS plus 0.05% tween. Cells were then blocked with 3% BSA and 0.3% triton in PBS for 1-2 hr. Following the blocking step, cells were treated with primary antibody overnight in blocking buffer. Primary antibodies used include anti-α and β-Synuclein (EP1646Y, N-terminal rabbit monoclonal, 1:100 dilution; Millipore, Billerica, MA), anti-pS129 αSyn (81A, mouse monoclonal, 1:1000 dilution, Biolegend; San Diego, CA), anti-pS129 αSyn (MBJR13, ab168381, Abcam, Cambridge, UK) and anti-MAP2 (ab5392, 1:10,000 dilution; Abcam, Cambridge, UK). Following incubation with antibodies, plates were washed 3-times with PBS containing 0.05% tween and then incubated 1 hr with fluorescent-conjugated secondary antibodies. The secondary antibodies used include Alexa Fluor 647 goat anti-mouse IgG (1:500 dilution); Alexa Fluor 488 goat anti-rabbit IgG (1:500 dilution), Alexa Fluor 568 goat anti-chicken IgG (1:500 dilution) and Hoechst, 1:800 dilution. Plates were then washed 3-times for 15 min each with PBS plus 0.05% tween, followed by a final wash in PBS alone.

### Immunofluorescent analysis

Images were acquired on ArrayScan™ VTi automated microscopy and image analysis system (Cellomics Inc., Pittsburgh, PA, USA) with a ×10 objective. Imaged plates were analyzed with the High Content Studio 3.0 software package using the neuronal profiling application. Cells were identified with Hoechst fluorescence which defines the nuclear area, and neurites were identified by MAP2 staining. The cell soma was identified by overlapping nuclear and MAP2 staining. The total insoluble αSyn and total insoluble pS129 αSyn were identified by the fluorescence intensities in two additional channels. Pathological induction is quantified by the total detergent-insoluble pS129 spot intensity co-localized to identified neurites. Fold induction is determined by normalizing to the mean of the negative control wells (untreated cells) within each plate. Z’ values were calculated for each plate based on negative control wells and 150 ng/ml PFF as the positive control. The average Z’ over the course of these experiments was 0.68. A toxicity index was calculated by multiplying the normalized nuclei count, normalized neuronal count, and normalized neurite length for each well. Values were normalized to the respective mean values of the negative control wells. Wells with toxicity scores of < 0.6 were excluded from analysis. Concentration response curves were generated and analyzed using Graphpad Prism (Graphpad Software, Inc. La Jolla, CA).

### Statistical analysis

Data expressed as average ± SD. Multiple comparisons were performed using one-way ANOVA or two-way ANOVA with Dunnett’s post hoc correction as appropriate.

## Acknowledgements

We are grateful to the Banner Sun Health Research Institute Brain and Body Donation Program of Sun City, Arizona and the Alzheimer's Disease Core Center, the Morris K. Udall Parkinson's Research Center of Excellence, and the Center for Neurodegenerative Disease Research, Perelman School of Medicine, the University of Pennsylvania (Philadelphia, PA) for providing human brain tissues. The Brain and Body Donation Program is supported by the National Institute of Neurological Disorders and Stroke (U24 NS072026 National Brain and Tissue Resource for Parkinson’s Disease and Related Disorders), the National Institute on Aging (P30 AG19610 Arizona Alzheimer’s Disease Core Center), the Arizona Department of Health Services (contract 211002, Arizona Alzheimer’s Research Center), the Arizona Biomedical Research Commission (contracts 4001, 0011, 05-901 and 1001 to the Arizona Parkinson’s disease Consortium) and the Michael J. Fox Foundation for Parkinson’s Research. The Alzheimer's Disease Core Center, the Morris K. Udall Parkinson's Research Center of Excellence, and the Center for Neurodegenerative Disease Research are supported by P30 AG10124 and P50 NS053488.

## Author Contributions

JDG, JEM, ND and MKA wrote the manuscript, designed the experiments and oversaw data generation and analysis. NH, CP, LY, LI and YC generated and analyzed the data and aided in the experimental design.

## References

1. Bendor JT, Logan TP, & Edwards RH (2013) The function of alpha-synuclein. Neuron 79(6):1044–1066.

2. Fauvet B, et al. (2012) alpha-Synuclein in central nervous system and from erythrocytes, mammalian cells, and Escherichia coli exists predominantly as disordered monomer. The Journal of biological chemistry 287(19):15345–15364.

3. Bartels T, Choi JG, & Selkoe DJ (2011) alpha-Synuclein occurs physiologically as a helically folded tetramer that resists aggregation. Nature 477(7362):107–110.

4. Wang W, et al. (2011) A soluble alpha-synuclein construct forms a dynamic tetramer. Proceedings of the National Academy of Sciences of the United States of America 108(43):17797–17802.

5. Beyer K & Ariza A (2013) alpha-Synuclein posttranslational modification and alternative splicing as a trigger for neurodegeneration. Mol Neurobiol 47(2):509–524.

6. Oueslati A (2016) Implication of Alpha-Synuclein Phosphorylation at S129 in Synucleinopathies: What Have We Learned in the Last Decade? Journal of Parkinson's disease 6(1):39–51.

7. Galvin JE, Lee VM, & Trojanowski JQ (2001) Synucleinopathies: clinical and pathological implications. Archives of neurology 58(2):186–190.

8. Chu Y & Kordower JH (2015) The prion hypothesis of Parkinson's disease. Current neurology and neuroscience reports 15(5):28.

9. Olanow CW & Brundin P (2013) Parkinson's disease and alpha synuclein: is Parkinson's disease a prion-like disorder? Movement disorders : official journal of the Movement Disorder Society 28(1):31–40.

10. Prusiner SB, et al. (2015) Evidence for alpha-synuclein prions causing multiple system atrophy in humans with parkinsonism. Proceedings of the National Academy of Sciences of the United States of America 112(38):E5308–5317.

11. Braak H, et al. (2003) Staging of brain pathology related to sporadic Parkinson's disease. Neurobiology of aging 24(2):197–211.

12. Kordower JH, Chu Y, Hauser RA, Freeman TB, & Olanow CW (2008) Lewy body-like pathology in long-term embryonic nigral transplants in Parkinson's disease. Nat Med 14(5):504–506.

13. Jin H, et al. (2008) Analyses of copy number and mRNA expression level of the alpha-synuclein gene in multiple system atrophy. Journal of medical and dental sciences 55(1):145–153.

14. Miller DW, et al. (2005) Absence of alpha-synuclein mRNA expression in normal and multiple system atrophy oligodendroglia. J Neural Transm (Vienna) 112(12):1613–1624.

15. Ozawa T, et al. (2001) Analysis of the expression level of alpha-synuclein mRNA using postmortem brain samples from pathologically confirmed cases of multiple system atrophy. Acta neuropathologica 102(2):188–190.

16. Asi YT, et al. (2014) Alpha-synuclein mRNA expression in oligodendrocytes in MSA. Glia 62(6):964–970.

17. Reyes JF, et al. (2014) Alpha-synuclein transfers from neurons to oligodendrocytes. Glia 62(3):387–398.

18. Watts JC, et al. (2013) Transmission of multiple system atrophy prions to transgenic mice. Proceedings of the National Academy of Sciences of the United States of America 110(48):19555–19560.

19. Polymeropoulos MH, et al. (1997) Mutation in the alpha-synuclein gene identified in families with Parkinson's disease. Science (New York, N.Y.) 276(5321):2045–2047.

20. Conway KA, Harper JD, & Lansbury PT (1998) Accelerated in vitro fibril formation by a mutant alpha-synuclein linked to early-onset Parkinson disease. Nature medicine 4(11):1318–1320.

21. Lee MK, et al. (2002) Human alpha-synuclein-harboring familial Parkinson's disease-linked Ala-53 --> Thr mutation causes neurodegenerative disease with alpha-synuclein aggregation in transgenic mice. Proceedings of the National Academy of Sciences of the United States of America 99(13):8968–8973.

22. Volpicelli-Daley LA, et al. (2011) Exogenous alpha-synuclein fibrils induce Lewy body pathology leading to synaptic dysfunction and neuron death. Neuron 72(1):57–71.

23. Tran HT, et al. (2014) Alpha-synuclein immunotherapy blocks uptake and templated propagation of misfolded alpha-synuclein and neurodegeneration. Cell reports 7(6):2054–2065.

24. Sacino AN, et al. (2014) Amyloidogenic alpha-synuclein seeds do not invariably induce rapid, widespread pathology in mice. Acta neuropathologica 127(5):645–665.

25. Uchihara T & Giasson BI (2016) Propagation of alpha-synuclein pathology: hypotheses, discoveries, and yet unresolved questions from experimental and human brain studies. Acta neuropathologica 131(1):49–73.

26. Koch JC, et al. (2015) Alpha-Synuclein affects neurite morphology, autophagy, vesicle transport and axonal degeneration in CNS neurons. Cell death & disease 6:e1811.

27. Woerman AL, et al. (2015) Propagation of prions causing synucleinopathies in cultured cells. Proceedings of the National Academy of Sciences of the United States of America 112(35):E4949–4958.

28. Takeda S, et al. (2015) Neuronal uptake and propagation of a rare phosphorylated high-molecular-weight tau derived from Alzheimer's disease brain. Nature communications 6:8490.

29. Foulds PG, et al. (2012) Post mortem cerebrospinal fluid alpha-synuclein levels are raised in multiple system atrophy and distinguish this from the other alpha-synucleinopathies, Parkinson's disease and Dementia with Lewy bodies. Neurobiology of disease 45(1):188–195.

30. Wong YC & Krainc D (2017) alpha-synuclein toxicity in neurodegeneration: mechanism and therapeutic strategies. Nat Med 23(2):1–13.

31. Stefanova N, Bucke P, Duerr S, & Wenning GK (2009) Multiple system atrophy: an update. The Lancet. Neurology 8(12):1172–1178.

32. Bousset L, et al. (2013) Structural and functional characterization of two alpha-synuclein strains. Nature communications 4:2575.

33. Ma MR, Hu ZW, Zhao YF, Chen YX, & Li YM (2016) Phosphorylation induces distinct alpha-synuclein strain formation. Scientific reports 6:37130.

34. Peelaerts W, et al. (2015) alpha-Synuclein strains cause distinct synucleinopathies after local and systemic administration. Nature 522(7556):340–344.

35. Raiss CC, et al. (2016) Functionally different alpha-synuclein inclusions yield insight into Parkinson's disease pathology. Scientific reports 6:23116.

36. Bae EJ, et al. (2012) Antibody-aided clearance of extracellular alpha-synuclein prevents cell-to-cell aggregate transmission. The Journal of neuroscience : the official journal of the Society for Neuroscience 32(39):13454–13469.

37. Games D, et al. (2014) Reducing C-terminal-truncated alpha-synuclein by immunotherapy attenuates neurodegeneration and propagation in Parkinson's disease-like models. The Journal of neuroscience : the official journal of the Society for Neuroscience 34(28):9441–9454.

38. Masliah E, et al. (2011) Passive immunization reduces behavioral and neuropathological deficits in an alpha-synuclein transgenic model of Lewy body disease. PloS one 6(4):e19338.

39. Sahin C, et al. (2017) Antibodies against the C-terminus of alpha-synuclein modulate its fibrillation. Biophysical chemistry 220:34–41.

40. Anderson JP, et al. (2006) Phosphorylation of Ser-129 is the dominant pathological modification of alpha-synuclein in familial and sporadic Lewy body disease. The Journal of biological chemistry 281(40):29739–29752.

41. Li W, et al. (2005) Aggregation promoting C-terminal truncation of alpha-synuclein is a normal cellular process and is enhanced by the familial Parkinson's disease-linked mutations. Proceedings of the National Academy of Sciences of the United States of America 102(6):2162–2167.

42. Prasad K, Beach TG, Hedreen J, & Richfield EK (2012) Critical role of truncated alpha-synuclein and aggregates in Parkinson's disease and incidental Lewy body disease. Brain pathology 22(6):811–825.

43. Michell AW, et al. (2007) The effect of truncated human alpha-synuclein (1-120) on dopaminergic cells in a transgenic mouse model of Parkinson's disease. Cell transplantation 16(5):461–474.

44. Ulusoy A, Febbraro F, Jensen PH, Kirik D, & Romero-Ramos M (2010) Co-expression of C-terminal truncated alpha-synuclein enhances full-length alpha-synuclein-induced pathology. The European journal of neuroscience 32(3):409–422.

45. Wakamatsu M, et al. (2008) Selective loss of nigral dopamine neurons induced by overexpression of truncated human alpha-synuclein in mice. Neurobiology of aging 29(4):574–585.

46. Bassil F, et al. (2016) Reducing C-terminal truncation mitigates synucleinopathy and neurodegeneration in a transgenic model of multiple system atrophy. Proceedings of the National Academy of Sciences of the United States of America 113(34):9593–9598.

47. Wang W, et al. (2016) Caspase-1 causes truncation and aggregation of the Parkinson's disease-associated protein alpha-synuclein. Proceedings of the National Academy of Sciences of the United States of America 113(34):9587–9592.

48. Fagerqvist T, et al. (2013) Monoclonal antibodies selective for alpha-synuclein oligomers/protofibrils recognize brain pathology in Lewy body disorders and alpha-synuclein transgenic mice with the disease-causing A30P mutation. Journal of neurochemistry 126(1):131–144.

49. Vaikath NN, et al. (2015) Generation and characterization of novel conformation-specific monoclonal antibodies for alpha-synuclein pathology. Neurobiology of disease 79:81–99.

50. Ishii R, et al. (2015) Decrease in plasma levels of alpha-synuclein is evident in patients with Parkinson's disease after elimination of heterophilic antibody interference. PloS one 10(4):e0123162.

51. Mollenhauer B, et al. (2011) alpha-Synuclein and tau concentrations in cerebrospinal fluid of patients presenting with parkinsonism: a cohort study. The Lancet. Neurology 10(3):230–240.

52. Unterberger U, et al. (2014) Detection of disease-associated alpha-synuclein in the cerebrospinal fluid: a feasibility study. Clinical neuropathology 33(5):329–334.

53. Tabrizi M, Bornstein GG, & Suria H (2010) Biodistribution mechanisms of therapeutic monoclonal antibodies in health and disease. The AAPS journal 12(1):33–43.

54. Mao X, et al. (2016) Pathological alpha-synuclein transmission initiated by binding lymphocyte-activation gene 3. Science 353(6307).

